# Increased CD33 levels tune activation and function of induced human microglial cells through inhibition of the TREM2 pathway

**DOI:** 10.64898/2026.01.28.701050

**Authors:** Amr Omer, Smita Jagtap, Elga Morrison, Lauren M. Carito, Alexander L. Prinzen, Stefan I. McDonough, Benjamin J. Andreone

## Abstract

The microglial *CD33* gene is overexpressed in the brains of individuals with Alzheimer’s disease, and multiple *CD33* genetic variants are directly linked to disease risk. Here, we investigate CD33 function in induced human microglial cells by increasing or decreasing CD33 levels with AAV6-mediated gene transfer and with CD33-targeting siRNA, respectively. Phagocytosis of oligomeric amyloid beta into live microglia was doubled by reducing CD33 levels, with concomitant increases in secreted TREM2 and in TREM2 activation as assayed by phosphorylation of downstream SYK. Increasing CD33 had opposing effects on microglial activity, decreasing amyloid beta uptake by approximately half, reducing LPS-induced increases in IL-10, and decreasing the efficacy of a TREM2-activating antibody. Experiments combining siRNA and AAV6 demonstrated a linear relationship between CD33 protein and amyloid beta uptake. Results show that CD33 largely governs multiple microglial functions associated with Alzheimer’s disease and affects both pharmacological and physiological activation of TREM2.

## Introduction

Microglia, the innate immune cells of the central nervous system (CNS), serve as resident phagocytes that dynamically survey the brain parenchyma, respond to acute injury, and protect against pathogens^1^. Activation and proliferation of microglia in the brain are prominent features of Alzheimer’s disease (AD), a neurodegenerative disorder marked by the accumulation of pathological amyloid plaques, neurofibrillary tangles, and neuronal cell death. In disease, activated microglia display beneficial functions, including increased phagocytosis, which is implicated in the clearance of amyloid plaques^2–4^. However, prolonged activation eventually contributes to microglial dysfunction, reduced phagocytosis, and the release of neurotoxic proinflammatory cytokines^5,6^.

Large-scale human genetic studies have associated AD pathogenesis with genes highly expressed in microglia^7–11^. Among these genes is *CD33,* which encodes a member of the sialic acid-binding immunoglobulin-like lectins (SIGLECs) family of transmembrane proteins expressed by immune and hematopoietic lineage cell types, including microglia. CD33 (also referred to as SIGLEC-3) signals through its immunoreceptor tyrosine-based inhibition motif (ITIM) domain to recruit the downstream phosphatases SHP-1/2 that dampen microglial activation. Mechanistically, CD33 acts upstream of the microglia-enriched transmembrane protein TREM2 to inhibit microglial activation and function. TREM2 signaling has been extensively implicated in beneficial microglial functions, including proliferation, amyloid plaque clearance, and the secretion of anti-inflammatory cytokines, including IL-10^12–15^. Upon ligand binding, TREM2 recruits the tyrosine kinase SYK to engage downstream microglial activation signaling cascades, including the AKT, MAPK, and JNK pathways that mediate microglial functionality. Besides acting as a surface receptor, TREM2 is also cleaved by ADAM proteases to generate secreted, soluble TREM2 (sTREM2)^16,17^. Although the precise function of sTREM2 is not fully understood, increased secretion of sTREM2 has been linked to TREM2 activation and is associated with improved disease prognosis^18,19^. Therefore, as mechanistic data suggest, activation and induced secretion of TREM2 via CD33 inhibition could serve as a promising therapeutic strategy to improve AD outcome.

Genome-wide association studies have linked SNPs in both the *TREM2* and *CD33* loci to AD risk. Carriers of the rs75932628 or rs143332484 SNPs within *TREM2* have an increased risk of AD^10,20^. These polymorphisms encode missense variants in TREM2 (R47H and R62H, respectively) that reduce ligand binding and subsequent signaling, indicating a loss of TREM2 function^21,22^. SNPs within the *CD33* locus have been associated with increased risk of (rs3865444C) and protection against (rs3865444A or rs12459419T) AD pathogenesis^23–26^. Protective SNPs within *CD33* are associated with increased expression of the minor isoform of CD33 protein lacking the extracellular ligand binding domain (CD33m), resulting in reduced expression of the major full-length CD33 isoform (CD33M). In contrast, the risk SNP within *CD33* is associated with increased expression of CD33M^27–29^. Together, these data support the opposing roles and functions of CD33 and TREM2 as regulators of microglial function in the context of AD.

Elevated full-length CD33 levels have been observed in the CNS of AD patients and are correlated with increased pathology, suggesting that reducing CD33 expression may offer protection against disease pathogenesis and progression^23,30–35^. In both mice and *in vitro* models, *CD33* knockout or downregulation is associated with increased amyloid clearance and decreased plaque burden^23,36–39^. However, despite the clear link between increased CD33 expression and AD pathology, previous models have focused on loss-of-function and have not recapitulated the increased CD33 levels seen in human disease. Therefore, a mechanistic understanding of the impact of CD33 overexpression on TREM2 signaling and secretion, amyloid clearance, and inflammation is not yet defined. It is also unknown whether CD33 downregulation has a beneficial effect on microglial function when CD33 levels are elevated, as in disease. Here, we model both CD33 overexpression and downregulation *in* vitro to demonstrate that CD33 levels are inversely correlated with TREM2 engagement and signaling, as well as microglial function. The data establish the inhibition of CD33 as a potential therapeutic approach to engage the TREM2 pathway in the treatment of AD.

## Results

### CD33 downregulation in microglia increases phagocytosis, TREM2 secretion, and TREM2 activation

To evaluate the impact of altered CD33 expression on microglial function, we used human induced pluripotent stem cell (iPSC)-derived microglia as a model system. iPSCs were differentiated into hematopoietic precursor cells (HPCs) and then into microglial-like cells (iMG) using established protocols^40^. Successful differentiation was validated at each stage by the expression levels of cell-type transcriptional markers, as well as upon iMG maturation by expression of the microglial-specific protein markers CX3CR1, IBA-1, and TMEM119 (Figures S1A-B).

Using this system, we first investigated the effect of CD33 downregulation using a divalent small interfering RNA (di-siRNA) targeting *CD33.* Di-siRNA is an emerging variant of siRNA technology in which each molecule contains two identical, covalently linked siRNA subunits that catalyze the cleavage of target mRNA transcripts via the RNA-induced silencing complex, resulting in subsequent protein downregulation. Di-siRNA has previously been shown to be taken up by neurons and microglia in vivo, inducing potent and durable target knockdown^41–45^. Two monovalent siRNAs targeting the *CD33* transcript sequence (NM_001772.4 in RefSeq) were transfected into iMG, resulting in a maximal 80% *CD33* knockdown at the highest concentration of 1nM, indicating a high level of activity (Figure S2). The more potent sequence (half-maximal inhibitory concentration of 1.95 pM) was synthesized as a di-siRNA for subsequent experiments, as this molecule will be appropriate for potential future experimentation in vivo.

To test whether CD33 knockdown by di-siRNA elicited an increase in microglial function, phagocytosis was assessed in iMG by measuring the uptake of amyloid beta. iMG were transfected with either 10 pM and 10 nM of di-siRNA for 24 hours and cellular uptake of pHrodo-labeled oligomerized amyloid beta was subsequently measured over a 20-hour window by live-cell imaging (Figure 1A). A fluorescent signal was detected only within phagocytic microglia, as pHrodo fluorescence is observed intracellularly within lower pH organelles, such as phagosomes, lysosomes, or late endosomes (Figure 1B). At both concentrations of di-siRNA, a nearly two-fold increase in amyloid beta uptake was observed by the end of the imaging window (Figure 1B-D). Knockdown of *CD33* transcript was confirmed by quantitative polymerase chain reaction in iMG cell lysates following imaging. Concentration-dependent reductions in mRNA levels were observed, with approximately 40% and 82% knockdown observed at 10 pM and 10 nM, respectively (Figure 1E).

**Figure 1:**
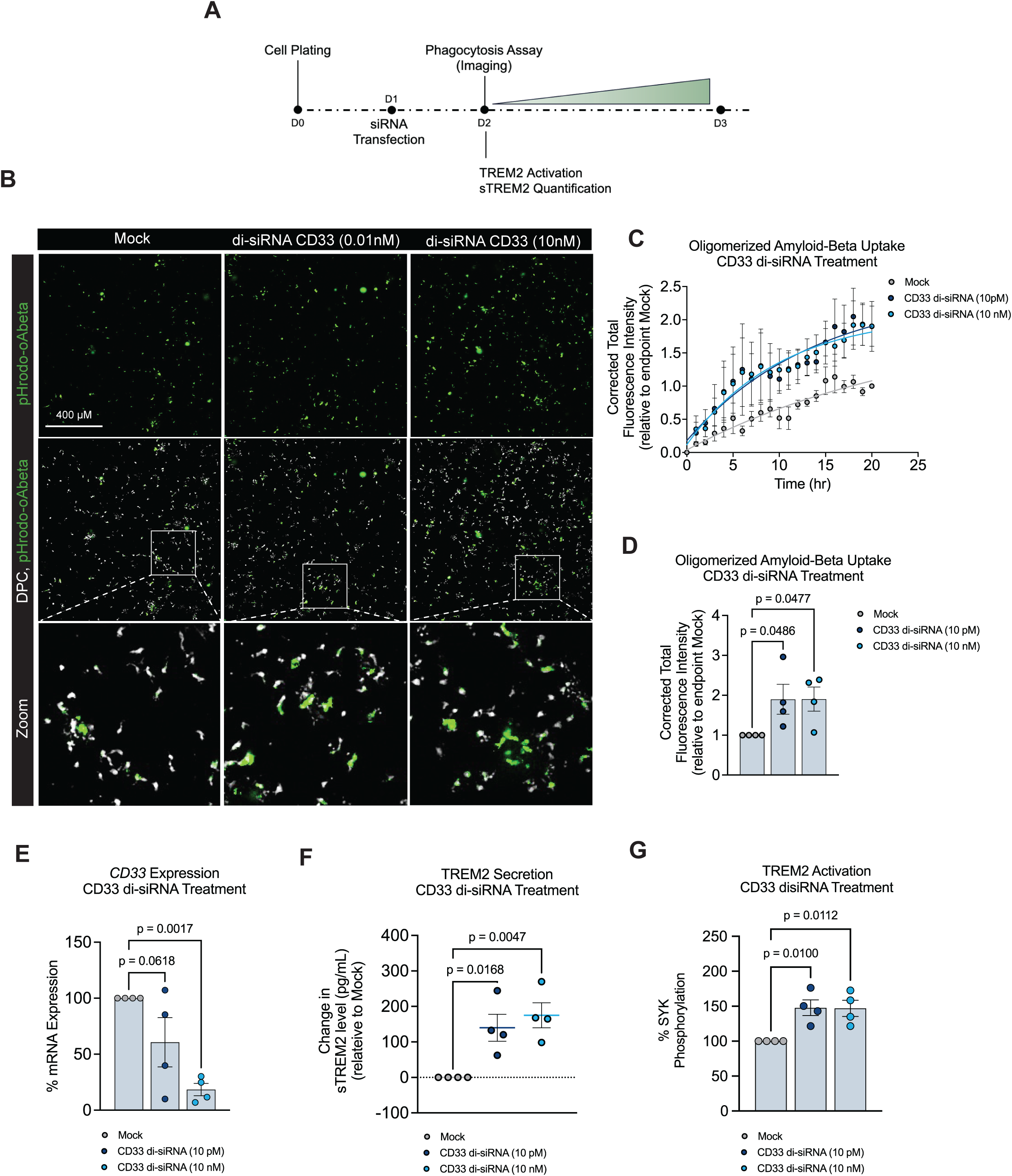
CD33 targeting di-siRNA increases phagocytosis of oligomerized amyloid beta, TREM2 secretion and activation in iMG. (A) Schematic of siRNA transfection of iMG. Cells were transfected 24 hours after iMG plating. After an additional 24 hours, cells were incubated with 2 μg/mL of pHrodo-labeled oligomerized amyloid (1-42) and imaged over a 20-hour window. (B) Confocal fluorescence images of iMG assessed for internalization of pHrodo-oligomerized amyloid beta treated as in (A). Digital phase contrast was used to visualize iMG. Images are stitched from four 20X images. Scale bar represents 400 μm. (C) Normalized intensity of internalized pHrodo-oligomerized amyloid beta in iMG per hour. Data are normalized to mock transfected control. (D) Normalized intensity of internalized pHrodo-oligomerized amyloid beta in iMG at endpoint. Data are normalized to mock transfected control. (E) RNA isolated from iMG 24 hours after transfection was used to quantify the percent *CD33* mRNA expression by qPCR. (F) Change in TREM2 secretion in conditioned media from iMG was quantified 24 hours after transfection by MSD. (G) Protein lysates isolated from iMG 24 hours after transfection were used to quantify percent SYK phosphorylation by AlphaLISA. Data represent mean ± SEM using one-way ANOVA with uncorrected Fisher’s LSD (D) and Dunnett’s multiple comparison test (E-G). Data points (n-numbers) are plotted on each bar graph and scatter plot each representing an independent experimental replicate. Line plots are representative of three independent experimental replicates (C).

To identify whether reductions in CD33 were linked to the TREM2 biology, we next evaluated TREM2 secretion by quantifying sTREM2 in the media of di-siRNA-treated iMG using an in-house developed MSD-based assay (Figure S3). Dose-dependent elevations in sTREM2 levels occurred following treatment, with approximately 140 and 175 pg/mL increases observed at 10 pM and 10 nM concentrations, respectively (Figure 1F). To identify whether reductions in CD33 were also linked to TREM2 activation, SYK phosphorylation was measured following di-siRNA treatment. At both 10 pM and 10 nM concentrations, an approximate 50% increase in phospho-SYK levels was observed (Figure 1G).

### CD33 overexpression in microglia impairs amyloid beta uptake and TREM2 secretion through reduced TREM2 expression

Given the large effect of CD33 knockdown on microglial phagocytosis, we investigated whether increasing levels of CD33, as observed in AD brain, also impacted amyloid beta uptake. We utilized adeno-associated virus (AAV) to overexpress full-length *CD33* (Ensembl ID: ENST00000262262). iMG were transduced with either a CD33-expressing AAV (AAV6-CD33) or a mCherry-expressing AAV control (AAV6-mCherry) for 72 hours and uptake of pHrodo-labeled oligomerized amyloid beta was measured for an additional 20 hours by live-cell imaging (Figure 2A). Transduction of iMG with AAV6-CD33 resulted in an approximately five-fold increase in CD33 protein levels compared to AAV6-mCherry or no AAV6 controls (Figure 2B). Elevated CD33 levels following AAV6-CD33 transduction led to a 50% decrease in the amyloid beta uptake by the end of the imaging window, compared to both AAV6-mCherry and no AAV6 controls (Figures 2C-E). Notably, transduction of iMG with increasing levels of AAV6-CD33 (0-25000 MOI) resulted in dose-dependent increases in CD33 protein expression and concomitant decreases in amyloid beta uptake (Figure S4A-B).

**Figure 2:**
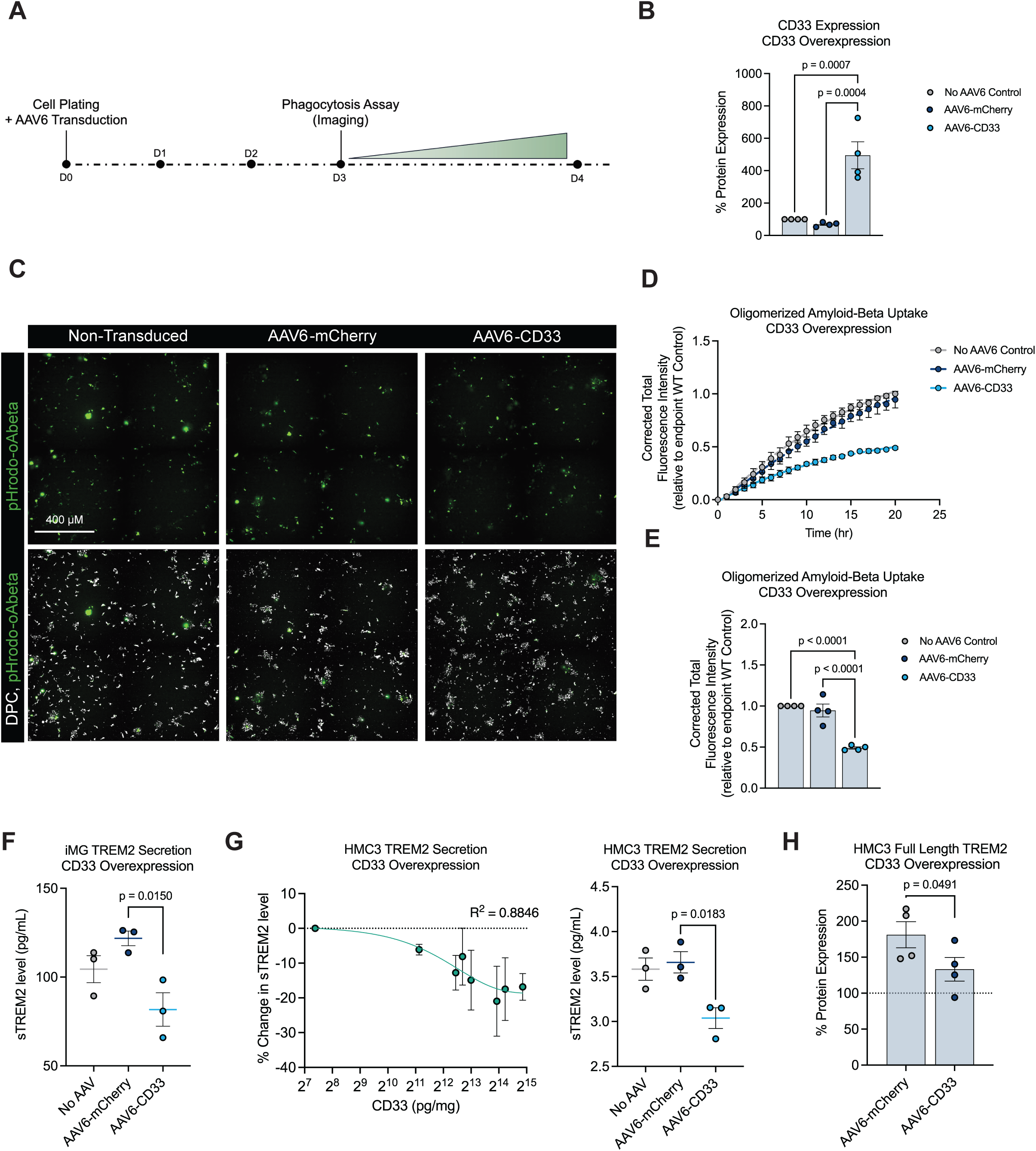
CD33 overexpression impairs phagocytosis of oligomerized amyloid beta and results in decreased TREM2 secretion in iMG and HMC3 cells. (A) Schematic of AAV6-mediated transduction of iMG. Cells were transduced with an MOI of 125000 at the time of plating. On day 3, cells were incubated with 2 μg/mL of pHrodo-labeled oligomerized amyloid (1-42) and imaged over a 20-hour window. (B) Protein lysates isolated from iMG 72 hours after transduction were used to quantify percent *CD33* protein expression by MSD. Data are normalized to no AAV6 transduction control. (C) Confocal fluorescence images of iMG assessed for internalization of pHrodo-oligomerized amyloid beta treated as in (A). Digital phase contrast was used to visualize iMG. Images are stitched from four 20X images. Scale bar represents 400 μm. (D) Normalized intensity of internalized pHrodo-oligomerized amyloid beta in iMG transduced with or without AAV6-CD33 or AAV6-mCherry per hour. Data are normalized to no AAV6 transduction control. (E) Normalized intensity of internalized pHrodo-oligomerized amyloid beta in iMG transduced with or without AAV6-CD33 or AAV6-mCherry at endpoint. Data are normalized to no AAV6 transduction control. (F) TREM2 secretion in conditioned media was quantified from iMG 72 hours after transduction by MSD. (G)(left) Percent change in TREM2 secretion in conditioned media was quantified from HMC3 cells 48 hours after transduction with increasing AAV6-CD33 concentrations by MSD. Data are represented as a percent change relative to dose-matched AAV6-mCherry. (right) TREM2 secretion in conditioned media was quantified from HMC3 cells 48 hours after transduction of AAV6-CD33 by MSD. (MOI = 250000). (H) Protein lysates isolated from HMC3 cells 48 hours after transduction were used to quantify percent full length TREM2 protein expression by MSD. (MOI = 250000). Data represent mean ± SEM using one-way ANOVA with post-hoc Tukey’s multiple comparisons test (B), with uncorrected Fisher’s LSD (E), with post-hoc Dunnett multiple comparisons test (F, G) and one-tailed unpaired t-test (H). Data points (n-numbers) are plotted on each bar graph and scatter plot each representing an independent experimental replicate. Line plots represent three independent experimental replicates (D, G).

To understand whether increases in CD33 protein modulated TREM2 cleavage, sTREM2 was quantified in the media of iMG 24 hours after AAV6-CD33 transduction. Although AAV6-mCherry transduction resulted in a 20% increase in sTREM2 levels compared to no AAV6 control, AAV6-CD33 transduction resulted in an approximately 40% reduction in soluble TREM2 levels (Figure 2F). We next sought to replicate these results in HMC3 human immortalized microglia and THP-1 human monocytes, as these models are readily amenable to the study of downstream signaling cascades. Similar to iMG, AAV6-CD33 transduction resulted in dose-dependent increases in CD33 protein levels in both immortalized cell models (Figure S5A-B). Corresponding dose-dependent decreases in sTREM2 levels also occurred, with approximately 10-20% reductions observed at the highest MOI evaluated in each cell line (Figure 2G and S5C). To understand the decrease in sTREM2 levels accompanying CD33 overexpression, full-length TREM2 levels were measured from HMC3 and THP-1 cell lysates. In concordance with the sTREM2 findings, AAV6-mCherry transduction augmented full-length TREM2 protein levels compared to no AAV6 control (Figure 2H and S5D). However, AAV6-CD33 transduction led to a 30-50% reduction in TREM2 protein in both HMC3 and THP-1 cells compared to AAV6-mCherry transduction, suggesting that CD33 negatively regulated sTREM2 secretion via reduction of full-length TREM2 (Figure 2H and S5D).

### CD33 overexpression impairs TREM2 activation and downstream anti-inflammatory signaling

As CD33 overexpression following AAV6-CD33 transduction reduced TREM2 expression, we tested whether TREM2 activation and signaling would also be inhibited (Figure 3A). We first measured baseline SYK phosphorylation in THP-1 cells following AAV6-CD33 transduction. An approximate 70% decrease in phospho-SYK levels was observed, relative to the AAV6-mCherry transduction (Figure 3B). To test whether this phenotype could be modulated by direct TREM2 engagement, THP-1 cells were transduced with AAV6-CD33 for 48 hours and subsequently treated with increasing concentrations of a TREM2 agonist antibody (Figure 3C). Phospho-SYK levels were lower in AAV6-CD33 transduced cells, with a 35% reduction compared to AAV6-mCherry transduction observed at the highest antibody concentration of 10 μg/mL, suggesting that direct TREM2 agonism was insufficient to overcome the inhibitory effect of CD33 on TREM2 activation (Figure 3C). In an orthogonal experiment, cells were transduced with increasing MOIs of AAV6-CD33 and subsequently treated with 2.5 µg/ml of TREM2 agonist antibody (Figure 3D). Dose-dependent reductions in phospho-SYK levels were observed following TREM2 antibody treatment, reaching saturation at an approximate 30% decrease in phospho-SYK, suggesting that TREM2 activation upon direct agonism is negatively regulated by CD33 (Figure 3D).

**Figure 3:**
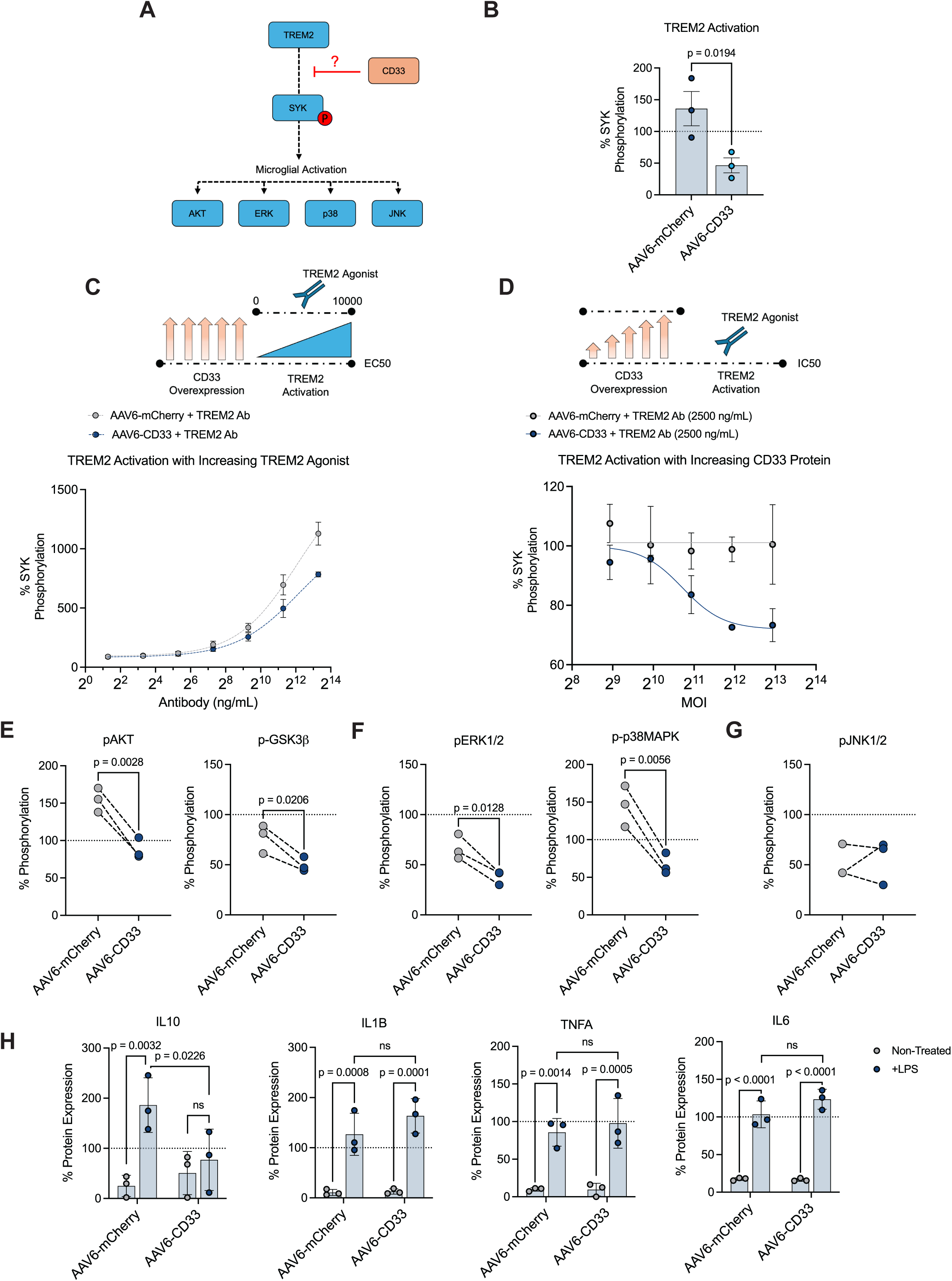
CD33 overexpression inhibits TREM2 activation, signaling, and anti-inflammatory secretion. (A) Schematic of the TREM2-CD33 pathway. CD33 inhibition acts on SYK activation, upstream of microglial activation pathways AKT, ERK, p38, and JNK. (B) Protein lysates isolated from THP-1 monocytes 48 hours after transduction were used to quantify percent SYK phosphorylation by MSD. (MOI = 31250) (C) 48 hours post-transduction, THP-1 cells (MOI = 31250) were treated with increasing concentrations of TREM2-targeting antibody agonist (0-10 μg/mL). After 5 minutes, protein lysates were isolated and used to quantify percent SYK phosphorylation by MSD. (D) 48-hours post-transduction of increasing concentrations of AAV6-CD33 (MOI = 0-31250), THP-1 cells were treated with a TREM2-targeting antibody agonist (2.5 ug/mL). After 5 minutes, protein lysates were isolated and used to quantify percent SYK phosphorylation by MSD. (E) Percent AKT and GSK3β phosphorylation was quantified from HMC3 cell lysates 48 hours after transduction of AAV6-CD33, quantified by MSD and ELISA, respectively. (F) Percent ERK and p38-MAPK phosphorylation was quantified from HMC3 cell lysates 48 hours after transduction of AAV6-CD33, quantified by ELISA. (G) Percent JNK phosphorylation was quantified from HMC3 cell lysates 48 hours after transduction of AAV6-CD33, quantified by ELISA. (H) (left to right) Percent expression of secreted IL10, IL1β, TNFα, and IL6 in conditioned media from HMC3 cells 24 hours after LPS challenge and 48 hours after transduction of AAV6-CD33 was quantified by MSD. Data represent mean ± SEM using one-tailed unpaired t-test (A, E-G), and two-way ANOVA using uncorrected Fisher’s LSD with a single pooled variance (H). Data points (n-numbers) are plotted on each bar graph and scatter plot each representing an independent experimental replicate. Line plots are representative of three and two independent experimental replicates (C, D, respectively).

We investigated further whether downstream microglial activation signaling pathways and the secretion of inflammatory cytokines were affected following CD33-dependent decreases in TREM2 activation and SYK phosphorylation. AKT signaling was inhibited following AAV6-CD33 transduction in HMC3 cells, with a 50% decrease in AKT phosphorylation and a 30% decrease in GSK3β phosphorylation observed (Figure 3E). A similar inhibition of the MAPK pathway occurred, with a 30% reduction in ERK1/2 phosphorylation and an 80% reduction in p38 MAPK phosphorylation observed (Figure 3F). The JNK pathway remained unaffected, with no observed change in JNK phosphorylation (Figure 3G). To test inflammation, HMC cells were challenged with LPS for 24 hours following AAV6-CD33 transduction and cytokine levels were measured in culture media. Baseline levels of each measured cytokine were unaffected by AAV6-CD33 transduction alone in the absence of LPS challenge (Figure 3H). While there were no significant changes in levels of the pro-inflammatory cytokines IL-1β, IL-6, and TNF-α after LPS challenge in AAV6-CD33 transduced cells, levels of the anti-inflammatory cytokine IL-10 were decreased by 54% in CD33 overexpressing cells compared to AAV6-mCherry transduced control cells following LPS challenge (Figure 3H).

### CD33-dependent impairment of phagocytosis is rescued with CD33 knockdown

Given that CD33 is elevated in AD brain and directly acts to impair TREM2-dependent microglial functionality, we tested whether CD33 knockdown could rescue phagocytosis when CD33 is overexpressed in iMG. 24 hours after transduction with either AAV6-CD33 or AAV6-mCherry control, iMG were treated with increasing concentrations of di-siRNA ranging from 0.001 nM to 0.1 nM. After a 48-hour treatment, uptake of pHrodo-labeled oligomerized amyloid beta was measured by live-cell imaging (Figure 4A). As previously observed (Figures 2C-E), AAV6-CD33 transduction led to a 50% decrease in the amyloid beta uptake by the end of the imaging window, compared to AAV6-mCherry control (Figures 4B-D). CD33 knockdown via di-siRNA resulted in dose-dependent restoration of amyloid beta uptake, with near complete rescue observed at the highest tested concentration of 0.1 nM (Figures 4B-D). Di-siRNA dose-dependent decreases in CD33 protein expression were also observed, with equivalent CD33 levels observed between the AAV6-CD33 group treated with 0.1 nM di-siRNA and the AAV6-mCherry control group (Figure 4E). Regression analysis further demonstrated a strong linear inverse correlation between levels of CD33 protein and amyloid beta uptake (Figure 4F).

**Figure 4:**
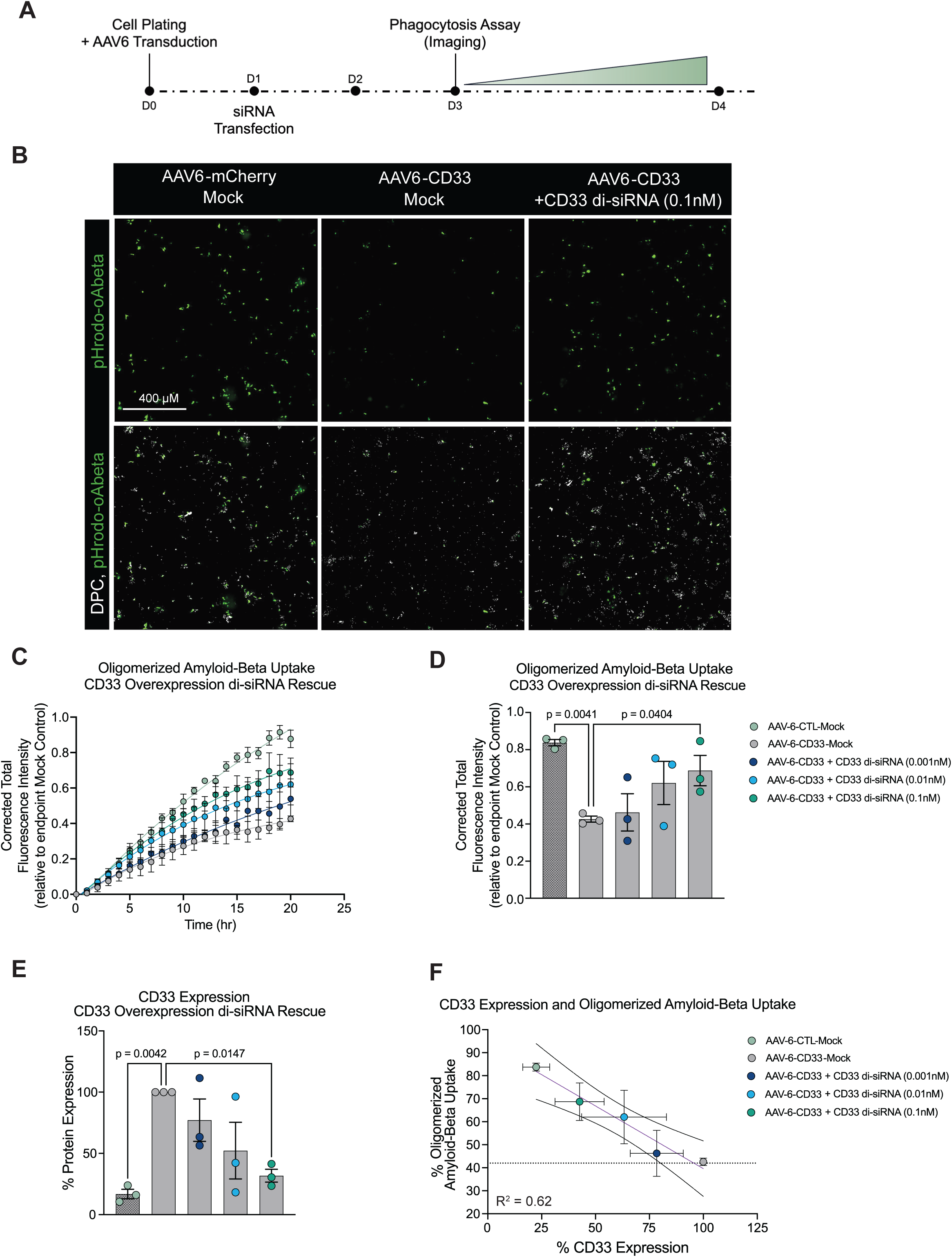
CD33 levels are inversely correlated with microglial phagocytosis in iMG. (A) Schematic of AAV6-mediated transduction of iMG. Cells were transduced with an MOI of 125000 at the time of plating. On day 1, cells were transfected with a di-siRNA targeting CD33 for 48 hours. On day 3, cells were incubated with 2 μg/mL of pHrodo-labeled oligomerized amyloid (1-42) and imaged over a 20-hour window. (B) Confocal fluorescence images of iMG assessed for internalization of pHrodo-oligomerized amyloid beta treated as in (A). Digital phase contrast was used to visualize iMG. Images are stitched from four 20X images. Scale bar represents 400 μm. (C) Normalized intensity of internalized pHrodo-oligomerized amyloid beta in iMG transduced with or without AAV6-CD33 or AAV6-mCherry per hour after transfection of increasing concentrations of CD33-targeting di-siRNA per hour. Data are normalized to mock transfected no AAV6 transduction control. (D) Normalized intensity of internalized pHrodo-oligomerized amyloid beta in iMG transduced with or without AAV6-CD33 or AAV6-mCherry per hour after transfection of increasing concentrations of CD33-targeting di-siRNA at endpoint. Data are normalized to mock transfected no AAV6 transduction control. (E) Protein lysates isolated from iMG 72 hours after transduction and 48 hours after transfection were used to quantify percent *CD33* protein expression by MSD. Data are normalized to mock transfected AAV6-CD33-treated iMG. (F) Linear regression of data from (D) and (E) was plotted to demonstrate the correlation between percent CD33 expression levels and oligomerized amyloid-beta uptake. Data represent mean ± SEM using one-way ANOVA with uncorrected Fisher’s LSD (D) and Dunnett’s multiple comparison test (E). Data points (numbers) are plotted on each bar graph and scatter plot, each representing an independent experimental replicate. Line plots are representative of three independent experimental replicates (C, F).

## Discussion

A mechanistic understanding of Alzheimer’s risk genes will likely prove important for developing AD therapeutics that leverage the innate immune system. While AD genetic risk variants in *CD33* result in increased expression^23,34^, the physiological effects of CD33 overexpression in human microglia have not yet been evaluated. Here, we show in induced human microglia, supported by experiments using monocyte-derived cell lines, that CD33 overexpression impairs microglial activation, likely by inhibiting TREM2 and its downstream signaling cascades. Data suggest that CD33 has a marked and internally consistent effect on multiple disease-relevant phenotypes and further support the role of CD33 as a critical player in AD pathogenesis. Our study builds on previous findings showing that additional AD-associated genes, such as *MS4A*, *PILRA*, and *LILRB2,* also negatively regulate TREM2 signaling and microglial function, underscoring the importance of the pathway as a whole^8,9,11^.

Bidirectional manipulation of CD33 levels produced large effects on phagocytosis, more than might be inferred from the small odds ratios of *CD33* genetic variants for Alzheimer’s risk^24,46^. Although AAV6 transduced only 30%-40% of iMG, the resulting CD33 overexpression reduced average amyloid beta uptake by 50%, an amount comparable to that seen in TREM2 knockout models *in vivo* and *in vitro*^47,48^. Reversal of this phenotype with di-siRNA was inversely proportional to CD33 protein levels and stoichiometric, suggesting that a broad range of CD33 expression may tune microglial function. Specifically, reducing CD33 levels in iMG boosted amyloid beta uptake by over 80%, exceeding previously described effects of TREM2 agonism^47,49^. Together, the large bidirectional effect of CD33 expression suggests that CD33 is capable of modulating the majority of TREM2-mediated phagocytosis. CD33 overexpression also significantly reduced the secretion of IL-10, which may increase AD risk by removing a counterbalance to pro-inflammatory factors such as IL-1β, IL-6, and TNF-α, the elevations of which are associated with Alzheimer’s disease through human data^50^.

Impaired TREM2 signaling capacity may be a key contributing factor to AD pathogenesis and the efficacy of TREM2-targeting therapeutics for disease treatment ^14^. Our findings demonstrate that CD33 overexpression reduced the capacity of a TREM2 agonist to activate TREM2 signaling, manifested as decreased phosphorylation of SYK and downstream effectors of the MAPK and AKT pathways. CD33 overexpression also reduced secreted sTREM2 levels, possibly due to inhibition of signaling pathways required to maintain full-length TREM2 expression ^51–53^. Consistent with the bidirectional effects of CD33 on phagocytosis, decreasing CD33 also produced increased sTREM2 levels. While the precise functions of sTREM2 in AD pathology are still under investigation, and the correlation of these in vitro measures to human physiology is unknown, increased sTREM2 concentrations in the cerebrospinal fluid of patients are associated with slower disease progression, including in subjects with autosomal-dominant Alzheimer’s ^16,17,54^. Therefore, for both sTREM2 and phagocytosis, the effects of decreasing CD33 were consistent with the protective genetic effect of *CD33* loss-of-function variants and with the beneficial effects of TREM2 activation and secretion. Since sTREM2 has emerged as a target engagement biomarker in clinical trials for molecules that directly activate the TREM2 protein ^55,56^, alterations in sTREM2 levels could also serve as a biomarker in the context of CD33 knockdown.

Given the crosstalk between CD33 and TREM2, targeted CD33 knockdown emerges as a novel strategy to activate TREM2 while retaining TREM2 activity and localization at the plasma membrane. The di-siRNA described here represents a promising approach to test the effects of CD33 knockdown in appropriate AD animal models, due to the potency, wide distribution, and months-long durability of di-siRNA in the brain ^41–43,45^. A CD33-targeting approach to AD may also have the potential to minimize amyloid-related imaging abnormalities (ARIA) and potential adverse consequences, likely caused by vascular complement activation in subsets of patients receiving high intravenous doses of anti-amyloid beta antibodies. The essentially normal physiology of adult humans missing CD33 entirely implies the safety of CD33 knockdown (although not necessarily in the context of AD).^57^ By enhancing endogenous microglial clearance, rather than directly mobilizing amyloid beta deposits with passive immunization, CD33-targeted therapies might circumvent ARIA while maintaining amyloid beta removal and possibly providing additional beneficial effects on sTREM2 and cytokine levels.

## Resource Availability

### Lead contact

Further information and requests for resources and reagents should be directed to the lead contact, Benjamin J. Andreone.

### Materials availability

All the original reagents presented in this study are available from the lead contact upon reasonable request.

### Data and code availability

All raw data generated in this study are presented in Table S1. All data reported in this paper will be shared by the lead contact upon request.

Any additional information required to reanalyze the data reported in this paper is available from the lead contact upon request.

## Author Contributions

A.O. performed experiments and produced figures. E.M. and S.J. performed iMG differentiation and IC50 assessment in iMGs. L.M.C. performed initial phospho-SYK assessments. A.P. led the synthesis of siRNA and di-siRNAs. A.O., B.J.A ., and S.I.M. wrote the manuscript with input from all other authors. A.O., S.J., S.I.M., and B.J.A. conceptualized and directed the execution of research goals.

## Supplemental information

**Figure S1: Validation of iMG differentiation**

(A) qPCR analysis of microglial differentiation panel comparing iPSC cells to HPCs iMG.

(B) JESS protein analysis of microglial-specific markers CX3CR1, IBA-1 and TMEM119.

**Figure S2: siRNAs targeting CD33 demonstrate high potency and dose-dependent knockdown of CD33 mRNA.**

RNA isolated from iMGs treated with independent monovalent siRNA at increasing concentrations for 24 hours were quantified using qPCR for determination IC50.

**Figure S3: Development of MSD assay to measure soluble TREM2 levels.**

Assay definition for TREM2 MSD used to assess total and soluble TREM2 levels.

**Figure S4: CD33-dependent inhibition of phagocytosis dose-dependent in iMG.**

(A) Protein lysates isolated from iMG 24 hours after transduction were used to quantify percent *CD33* protein expression by MSD. Data are normalized to no AAV6 transduction controls.

(B) Normalized intensity of internalized pHrodo-oligomerized amyloid beta in iMG transduced with or without AAV6-CD33 or AAV6-mCherry at endpoint. Data are normalized to no AAV6 transduction control.

Data represent mean ± SEM. Data points (numbers) are plotted on each bar graph, each representing an independent technical replicate.

**Figure S5: CD33 overexpression impairs TREM2 secretion in THP-1 cells.**

(A) Protein lysates isolated from HMC3 cells 48 hours after transduction of increasing concentrations of AAV6-CD33 were used to quantify percent *CD33* protein expression by MSD.

(B) Protein lysates isolated from THP-1 cells 48 hours after transduction of increasing concentrations of AAV6-CD33 were used to quantify percent *CD33* protein expression by MSD.

(C) (left) Percent change in TREM2 secretion in conditioned media was quantified from THP-1 cells 48 hours after transduction with increased AAV6-CD33 concentrations by MSD. Data are represented as a percent change relative to dose-matched AAV6-mCherry. (right) TREM2 secretion in conditioned media was quantified from THP-1 cells 48 hours after transduction of AAV6-CD33 by MSD. (MOI = 31250).

(D) Protein lysates isolated from HMC3 cells 48 hours after transduction were used to quantify percent full length TREM2 protein expression by MSD. (MOI = 31250).

Data represent mean ± SEM using one-way ANOVA with post-hoc Dunnett multiple comparisons test (C) and one-tailed unpaired t-test (D). Data points (n-numbers) are plotted on each bar graph and scatter plot each representing an independent experimental replicate. Line plots represent three independent experimental replicates (C).

**Figure S6: AAV6 serotype produces the highest transduction efficiency in iMG.**

(A) Confocal fluorescence images of iMG 24 hours after transduction of AAV2, 6, 8, 9, and PHP.eB expressing EGFP. Digital phase contrast was used to visualize iMG. Images are stitched from four 20X images. Scale bar represents 400 μm.

(B) Quantification of percentage of cells positive for EGFP expression with increasing AAV dosage (left) 24 hours and (right) 48 hours post-transduction.

(C) Quantification of percentage of cells positive for EGFP expression over time after AAV6 transduction at indicated MOIs.

(D) Quantification of EGFP mean intensity of cells over time after AAV6 transduction at indicated MOIs.

Data represent mean ± SEM. Line plots represent three technical replicates (B-D).

## STAR Methods

### EXPERIMENTAL MODEL AND STUDY PARTICIPANT DETAILS

#### iPSC maintenance and microglial differentiation

iPSC Maintenance: iPSCs were purchased from ATTC (ATCC-BXS0115) and cultured in mTeSR Plus media (STEMCELL Technologies) in Matrigel-coated 6-well plates (Corning), with subculturing every five to seven days, depending on confluency.

Hematopoietic differentiation: Differentiation of iPSCs into iPSC-derived hematopoietic progenitor cells (iHPCs) was performed using the STEMdiff Hematopoietic kit (STEMCELL Technologies). On day -1, cells were detached into small colony aggregates measuring 100-200 µm in diameter using the StemPro EZPassage™ Disposable Stem Cell Passaging Tool (Invitrogen). About 40-80 aggregates were then plated in Matrigel-coated 12-well plates with 1 mL of mTeSR Plus yielding 16-40 colonies per well for differentiation. On day 0, 1 mL of STEMdiff hematopoietic medium A was added. On day 2, half of the cell supernatants were replaced with fresh medium A. On day 3, the medium was fully replaced with 1 mL of STEMdiff hematopoietic medium B. On days 5, 7, and 10, half of the supernatants were replaced with fresh medium B. On day 12, the floating iHPCs in the cell supernatants were collected and centrifuged at 300 g for 5 minutes. Each harvest is expected to yield between 0.1 and 1 × 10^6^ iHPCs per well.

Microglial differentiation: On day 0, iHPCs were resuspended at a density of 5 × 10^5^ cells/mL in microglia differentiation medium and plated on Matrigel-coated 6-well plates. The culture was supplemented with 1 mL of differentiation medium per well every other day. Every 10 to 12 days, 5 mL of cell supernatants were collected from each well and centrifuged at 300×g for 5 minutes. The collected cells were then resuspended in 1 mL of differentiation medium and returned to culture. Cells were regarded as mature iMG on day 28-34 of microglial differentiation. The iMG were cryopreserved in liquid nitrogen using mFreSR at a concentration of 1×10^6^ per vial.

#### Immortalized cell culture

HMC3 Cells: The human microglial cell line HMC3 (CRL-3304) was obtained from ATCC. Cells were cultured in EMEM medium supplemented with 10% FBS (Gibco) and kept in a 37 °C incubator with 5% CO2.

THP-1 Cells: The human monocytic cell line THP-1 (TIB-202) was obtained from ATCC. Cells were cultured in RPMI medium supplemented with 10% FBS (Gibco) and kept in a 37 °C incubator with 5% CO2.

### METHOD DETAILS

#### siRNA and di-siRNA synthesis

Monovalent and divalent siRNAs were synthesized in-house using modified (2’-F, 2’-OMe) phosphoramidites with standard protecting groups on solid support as described previously^41^. siRNAs contained a 21-mer antisense strand and a 16-mer sense strand, with the antisense strand targeting the *CD33* transcript sequence.

Sense and antisense single strands of monovalent siRNA were synthesized at 1 µmole scale using Dr. Oligo 48 (Biolytic). Sense strand was synthesized using cholesterol-conjugated CPG (500A, LGC Biosearch Technologies), while antisense strand was synthesized using Unylinker® CPG (500A, LGC Biosearch Technologies). Mid-scale di-siRNA synthesis was performed using MerMade 12 (LGC Biosearch Technologies) or AKTA oligopilot 10 (Cytiva). Divalent oligonucleotide sense strand was grown on modified solid support custom-synthesized by Hongene Biotech as originally described^41^. Antisense strand containing 5’- vinylphosphonate (Vp) was synthesized on Unylinker CPG. Standard oligonucleotide synthesis reagents were purchased from ChemGenes. Modified phosphoramidites (Hongene Biotech), including 2’-F, 2’-Ome, 5’-Vp and 5’-Phosphate with standard protecting group, were prepared at 0.1 M for Dr. Oligo 48 and MerMade, and 0.2 M for AKTA oligopilot 10. All oligonucleotides were synthesized as reported with minor updates as follows: Cap B reagent was a mixture of 1:1 by volume of 40% Acetic anhydride in ACN (RN-2217, ChemGenes) and 60% symmetrical collidine in ACN (RN-2225, ChemGenes).

Small-scale synthesis was deprotected and cleaved from the solid support under methylamine gas at room temperature for 50 min. Mid-scale synthesis followed a regular deprotection and cleavage path using concentrated liquid ammonium hydroxide at 55°C for 18 hours. Purifications on IEX HPLC were performed at the single-strand level. Sense and antisense strands were then annealed in 1xPBS at 95°C for 10 min followed by slow cooling at room temperature for 2 hours. IEX HPLC confirmed both single-strand and duplex purity, and the identity of the oligonucleotide was confirmed using ESI-LC/MS. The di-siRNA was formulated in 1X PBS.

#### AAV production, selection, and transduction

AAV serotyping in iMG was completed using VectorBuilder *in vivo* grade AAV serotype testing panel (CMV-EGFP). The transduction efficiency of enhanced green fluorescent protein (EGFP)-expressing AAVs of five distinct serotypes from the AAV serotype testing panel (AAV2, 6, 8, 9, and PHP.eB) was tested using live cell imaging. Each AAV used a cytomegalovirus (CMV) promoter followed by enhanced green fluorescent protein (EGFP), thereby enabling robust expression of EGFP for live-cell fluorescence imaging of iMG across a broad range of viral MOIs. AAVs were diluted to the desired concentration in Opti-MEM (Gibco). 100 μL of diluted AAV was added to the wells. Subsequently, 100 μL iMG were plated at 3 × 10^4^ cells per well in PDL-coated 96-well plates (Phenoplate, Revvity). Cells were incubated in a 37 °C incubator with 5% CO2.

AAV6 was selected, as it exhibited the highest level of transduction, with approximately 20-30% of cells expressing EGFP after 72 hours at a saturating multiplicity of infection (MOI) of 1.2×10^5^ (Figures S6A-B). In the case of AAV6, the imaging window was extended to 144 hours to evaluate the influence of viral dosage on EGFP expression and intensity. The data indicate that optimal conditions for overexpression occur between 48 and 72 hours (Figure S6C-D).

After 48 hours, live cell imaging was conducted using the Operetta CLS High-Content Analysis System. Imaging was done at 5% CO_2_ and 37 °C at 1-hour intervals. Individual cells were identified using digital phase contrast channel to discriminate between microglia. EGFP intensity and positivity was determined on a per-cell basis using the Harmony HCA Software (Revvity).

VectorBuilder produced AAV6-CD33 using CD33 (NM_001772.4). A CMV promoter drove CD33 overexpression. AAV6-mCherry was obtained from Charles River Laboratories. For transduction, AAVs were diluted to the desired concentration in Opti-MEM (Gibco). 100 μL of diluted AAV was added to the wells. Subsequently, 100 μL of cells were plated at 3 × 10^4^ cells per well in PDL-coated 96-well plates (Phenoplate, Revvity). After 24 hours, media was exchanged. After an additional 24 hours, cells were processed for downstream experimental assessment.

#### RNAiMAX-based siRNA and di-siRNA transfection

siRNAs or di-siRNAs were actively transfected after complexing in 0.5% RNAiMAX as per the manufacturer’s protocol.

For IC50 determination in iMG, cells were plated in Corning 96-well plates. After 24 hours, cells were transfected with siRNAs at concentrations ranging from 100 nM to 1 fM for 48 hours.

For the remaining experiments, iMG were plated in PDL-coated Corning 96-well plates. After 24 hours, cells were transfected with di-siRNA for an additional 24 hours for downstream functional assessment.

#### Glial-Mag-based di-siRNA transfection

For CD33 knockdown after AAV transduction, di-siRNA was actively transfected after complexing in 0.2% Glial-Mag as per the manufacturer’s protocol 24 hours post AAV transduction. Subsequent phagocytic assessment or protein measurements were completed 24 hours after transfection.

#### Real-time qPCR and *CD33* transcript quantification

RNA was isolated using PureLink RNA Mini Kit (Invitrogen) as per the manufacturer’s protocol. RT-qPCR was performed using the TaqMan Fast Advanced Cells-to-Ct Kit (ThermoFisher). cDNA was synthesized using the High-Capacity cDNA RT Kit (Applied Biosystems) as per the manufacturer’s protocol. CD33 mRNA expression was compared to mock-transfected cells using the ΔΔCt method.

cDNA samples underwent qPCR analysis using TaqMan Fast Virus 1-Step (Invitrogen) and probes (*ATP5b* housekeeping gene: Hs00969569_m1, *CD33*: Hs00233544_m1) on a QuantStudio 7 Flex.

#### CD33 protein quantification

CD33 protein levels were measured using CD33 R-PLEX assay (Meso Scale Diagnostics, Rockville, MD). Cells were lysed in 50 μL of 1X RIPA Buffer (EMD Millipore). In HMC3 and THP-1 cells, lysates were isolated 48 hours post-transduction. In iMGs, lysates were isolated 72 hours post-transduction. Samples were analyzed as per the manufacturer’s protocol. The concentration of CD33 was interpolated using calibrators on MSD Discovery Workbench analysis software.

#### pHrodo oligomerized amyloid beta (1-42) phagocytosis assay and live-cell imaging

Oligomerized amyloid-beta (1-42) particles (StressMarg) were obtained and pHrodo-labeled using pHrodo iFL Green STP ester labeling kit (Invitrogen), following the manufacturer’s instructions.

For phagocytosis assays, iMG were plated at 3 × 10^4^ cells per well in PDL-coated 96-well plates (Phenoplate, Revvity). Non-transduced iMGs were assessed 24 hours post-transfection with di-siRNA. Transduced iMGs were assessed 72 hours post-transduction, corresponding to 48-hours post-transfection. Post-treatment, 2 μg/mL of pHrodo-labeled oligomers were added per well, and cells were subsequently imaged for 20 hours.

Live cell imaging was conducted using the Operetta CLS High-Content Analysis System. Imaging was done at 5% CO_2_ and 37 °C at 1-hour intervals. Individual cells were identified using digital phase contrast channel to discriminate between microglia. pHrodo intensity was determined on a per-cell basis through measurement of total fluorescence using the Harmony HCA Software (Revvity).

#### Total and soluble TREM2 protein quantification

A custom human TREM2 MSD-based assay was developed to measure TREM2 expression or sTREM2 in conditioned media. In HMC3 and THP-1 cells, conditioned media were collected 48 hours post-transduction. In iMGs, conditioned media was collected 24 hours post-transfection. Briefly, streptavidin-coated small spot MSD plates (Meso Scale Diagnostics, Rockville, MD) were coated with Biotinylated goat anti-human TREM2 polyclonal antibody (R&D Systems) used as a capture antibody for 1.5 hours with agitation at 600 rpm. Plates were washed three times in 1X PBS-Tween (ThermoFisher). 25 μL of sample and TREM2 calibrators (R&D Systems) were added and incubated for 1 hour at room temperature with agitation at 600 rpm. Plates were washed three times in 1X PBS-Tween (ThermoFisher). SULFO-TAG labeled TREM2 Ab (Abcam) was added as detection antibody and incubated at room temperature with agitation at 600 rpm. Plates were washed three times in 1X PBS-Tween (ThermoFisher). The detection antibody was labeled with SULFO-TAG according to the manufacturer’s instructions (Meso Scale Diagnostics, Rockville, MD). Plates were washed three times in 1X PBS-Tween (ThermoFisher). 150 uL of MSD Gold Read Buffer (Meso Scale Diagnostics, Rockville, MD) was added to each well and plates were analyzed on a Sector Imager (Meso Scale Diagnostics, Rockville, MD). Concentrations of sTREM2 or TREM2 were interpolated using calibrators on MSD Discovery Workbench analysis software.

#### TREM2 activation measured by phospho-SYK AlphaLISA

24 hours after iMG transfection, iMG lysates were assayed using the standard protocol for a commercial AlphaLISA pSyk assay (PerkinElmer). Ten microliters of lysate per well were transferred to a white opaque 384-well OptiPlate (PerkinElmer). Next, 5 μL of Acceptor Mix was added per well and incubated for 1 h at room temperature. Then, 5 μL of Donor Mix was added per well under reduced light conditions. Plates were again incubated for 1 h at room temperature and read using AlphaLISA settings on a PerkinElmer EnVision plate reader.

#### TREM2 activation measured by phospho-SYK MSD

A custom human phospho-SYK MSD-based assay was developed to measure phosphorylated and total SYK expression in THP-1 cells. Briefly, 48 hours after AAV6 transduction, THP-1 cell samples were lysed in 50 μL of 1X RIPA Buffer (EMD Millipore). Streptavidin-coated small spot MSD plates (Meso Scale Diagnostics, Rockville, MD) were coated with Biotinylated goat anti-human SYK or phospho-SYK monoclonal antibody (Cell Signaling Technology) used as a capture antibody for 1 hour with agitation at 600 rpm. Plates were washed three times in 1X PBS-Tween (Thermofisher). 25 μL of sample was added and incubated for 1 hour at room temperature with agitation at 600 rpm. Plates were washed three times in 1X PBS-Tween (Thermofisher). SULFO-TAG labeled total-SYK monoclonal antibody (Cell Signaling Technology) was added as detection antibody and incubated at room temperature with agitation at 600 rpm. Plates were washed three times in 1X PBS-Tween (Thermofisher). Detection antibody was labeled with SULFO-TAG as per manufacturer’s instructions (Meso Scale Diagnostics, Rockville, MD). Plates were washed three times in 1X PBS-Tween (Thermofisher). 150 uL of MSD Gold Read Buffer (Meso Scale Diagnostics, Rockville, MD) was added to each well and plates were analyzed on a Sector Imager (Meso Scale Diagnostics, Rockville, MD). Raw signals of phospho-SYK or total SYK were used to determine normalized percent changes compared to controls.

#### AKT and MAPK activation panels

HMC3 cells were lysed in 1X RIPA Buffer (EMD Millipore) 48 hours post-transduction with AAV6-CD33. Lysates were quantified using the Pierce BCA assay (ThermoFisher), and total protein concentrations, alongside BSA calibrators (ThermoFisher), were measured on a Spectramax plate reader using SoftMax Pro software to determine total protein concentrations.

Phospho-AKT levels were measured by MSD using T-PLEX assay (Meso Scale Diagnostics, Rockville, MD) as per the manufacturer’s protocol using HMC3 cell lysates. Raw signals were normalized against their corresponding total protein concentrations.

Phospho-p38 MAPK, phospho-ERK1/2, and phospho-JNK1/2/3 levels were measured by ELISA using the multispecies MAPK Family Activation InstantOne ELISA Kit (Invitrogen) as per the manufacturer’s protocol using HMC3 cell lysates. Raw signals were normalized against their corresponding total protein concentrations.

Phospho-GSK3β levels were measured by ELISA using PathScan Phospho-GSK-3β (Ser9) Sandwich ELISA Kit (Cell Signaling Technology) as per the manufacturer’s protocol using HMC3 cell lysates. Raw signals were normalized against their corresponding total protein concentrations.

All normalized signals were corrected against experimental controls to determine the percent change of phosphorylation.

#### LPS inflammatory challenge

48 hours after AAV transduction, HMC3 cells were treated with 100 ng/mL of LPS (Invitrogen) for 24 hours. Conditioned media were collected, and the secretion of inflammatory factors was assessed by MSD pro-inflammatory cytokine panel.

#### MSD pro-inflammatory cytokine panel

Inflammatory factor secretion was measured using the R-PLEX IL-10 assay and the 4-PLEX V-PLEX pro-inflammatory panel II kit (Meso Scale Diagnostics, Rockville, MD). HMC3 conditioned media were analyzed 48 hours after AAV transduction as per the manufacturer’s protocol. Concentrations of pro-inflammatory factors were interpolated using calibrators on MSD Discovery Workbench analysis software.

#### TREM2 antibody-dependent agonism

Human TREM2 agonistic antibody was acquired from R&D Systems. THP-1 cells were transduced with AAV6-CD33, and after 48 hours, the cells were treated with 2.5 μg/mL of TREM2 antibody for IC50 assessment or with increasing concentrations (0-10 μg/mL) of TREM2 antibody for EC50 assessment for 5 minutes. Cells were lysed in 1X RIPA Buffer (EMDMillipore). pSYK and SYK were measured by MSD as described above.

### QUANTIFICATION AND STATISTICAL ANALYSIS

All statistical analyses were performed using Prism (GraphPad). All statistical analyses are described within the figure legends. Two-tailed Student’s *t* tests were used to determine statistically significant differences between two groups. One-way ANOVA or two-way ANOVA was used for multiple comparisons, and post-hoc tests are described in the figure legends. Nonlinear regression was performed and is described in Table S1. R-squared values are displayed. *P*-values are displayed on all bar graphs where applicable. For ANOVAs, *F* values and degrees of freedom are provided in Table S1. For *t* tests, *t* values and degrees of freedom are provided in Table S1.

