## Supplementary figures and images for "Increased CD33 levels tune activation and function of induced human microglial cells through inhibition of the TREM2 pathway"

### Supplemental Figure

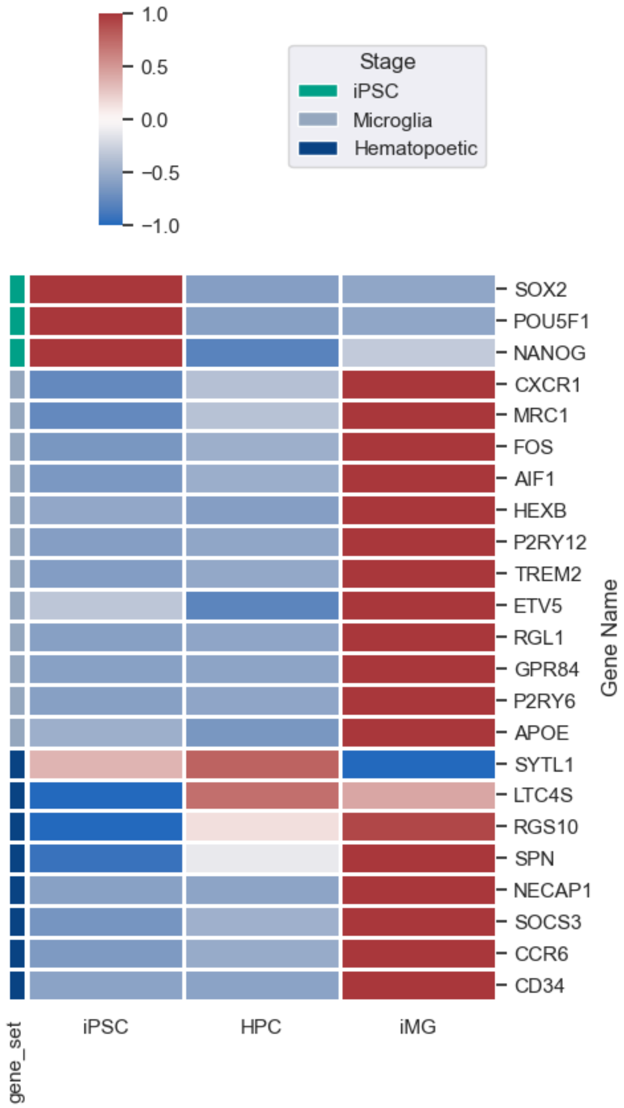

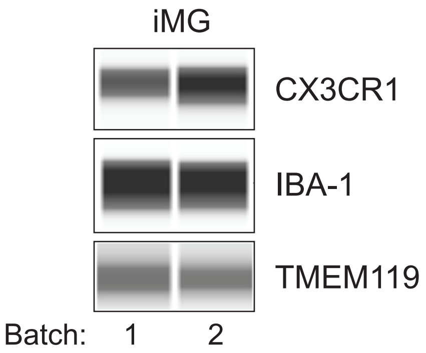

#### Figure S2

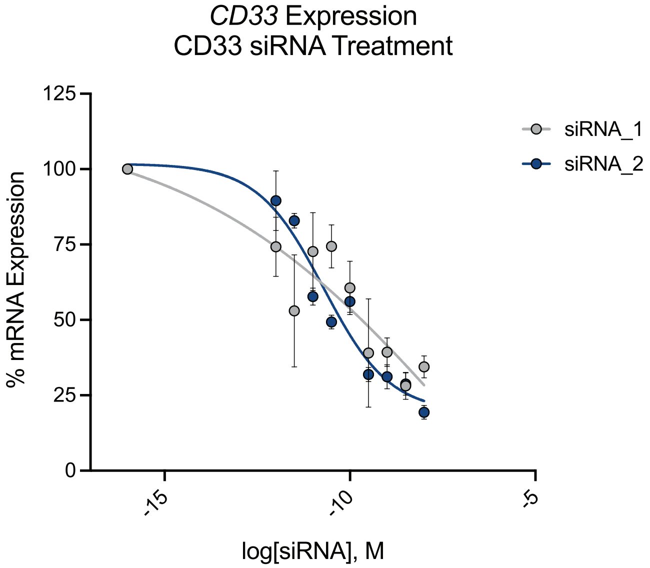


| | siRNA_1 | siRNA_2 |
|---|---|---|
| IC50 | 1.359e-007 | 1.949e-011 |

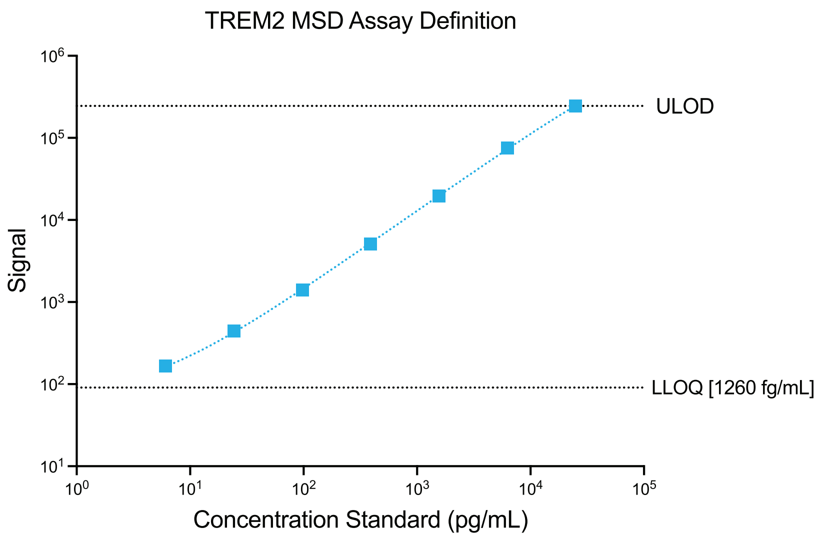

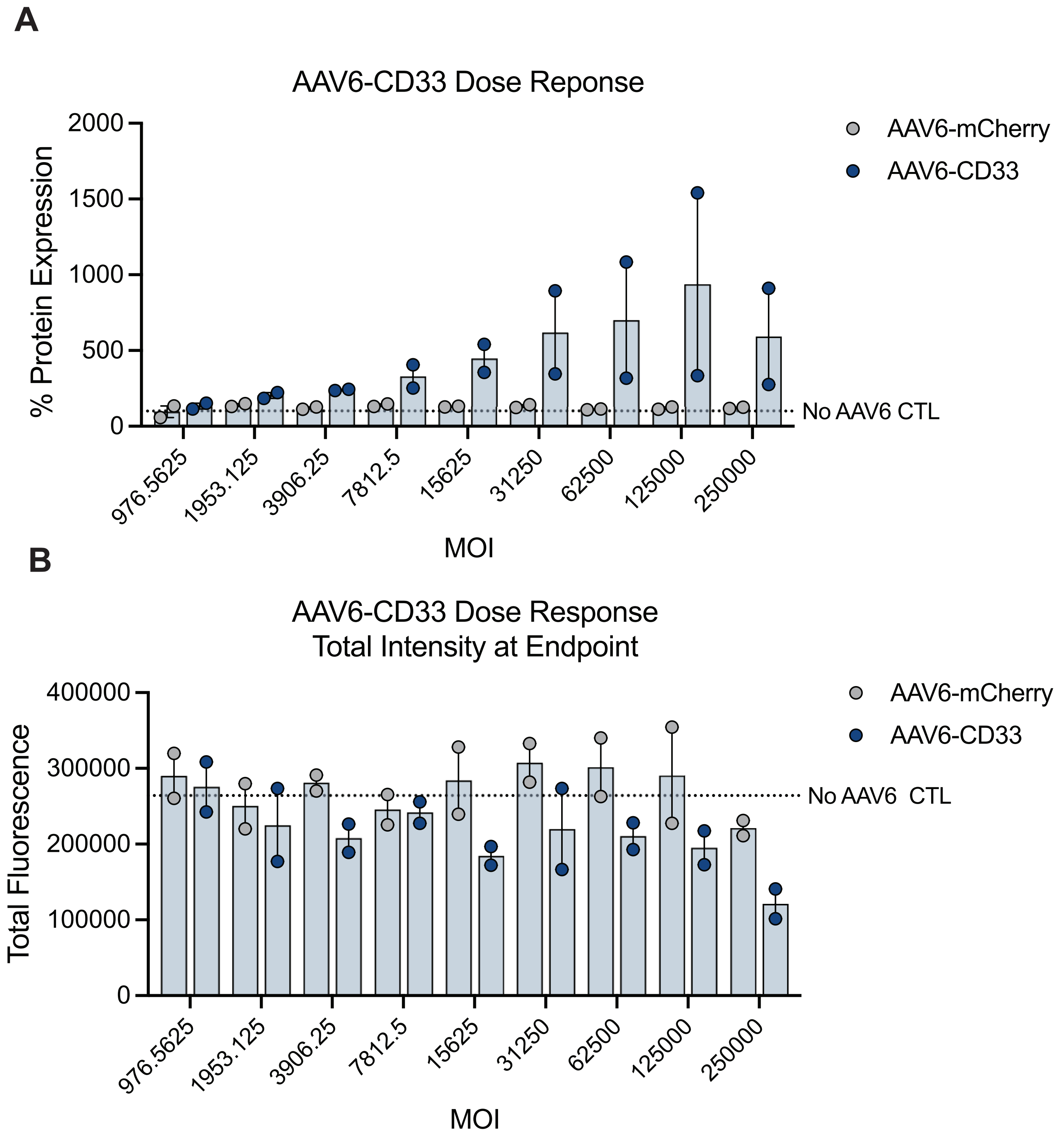

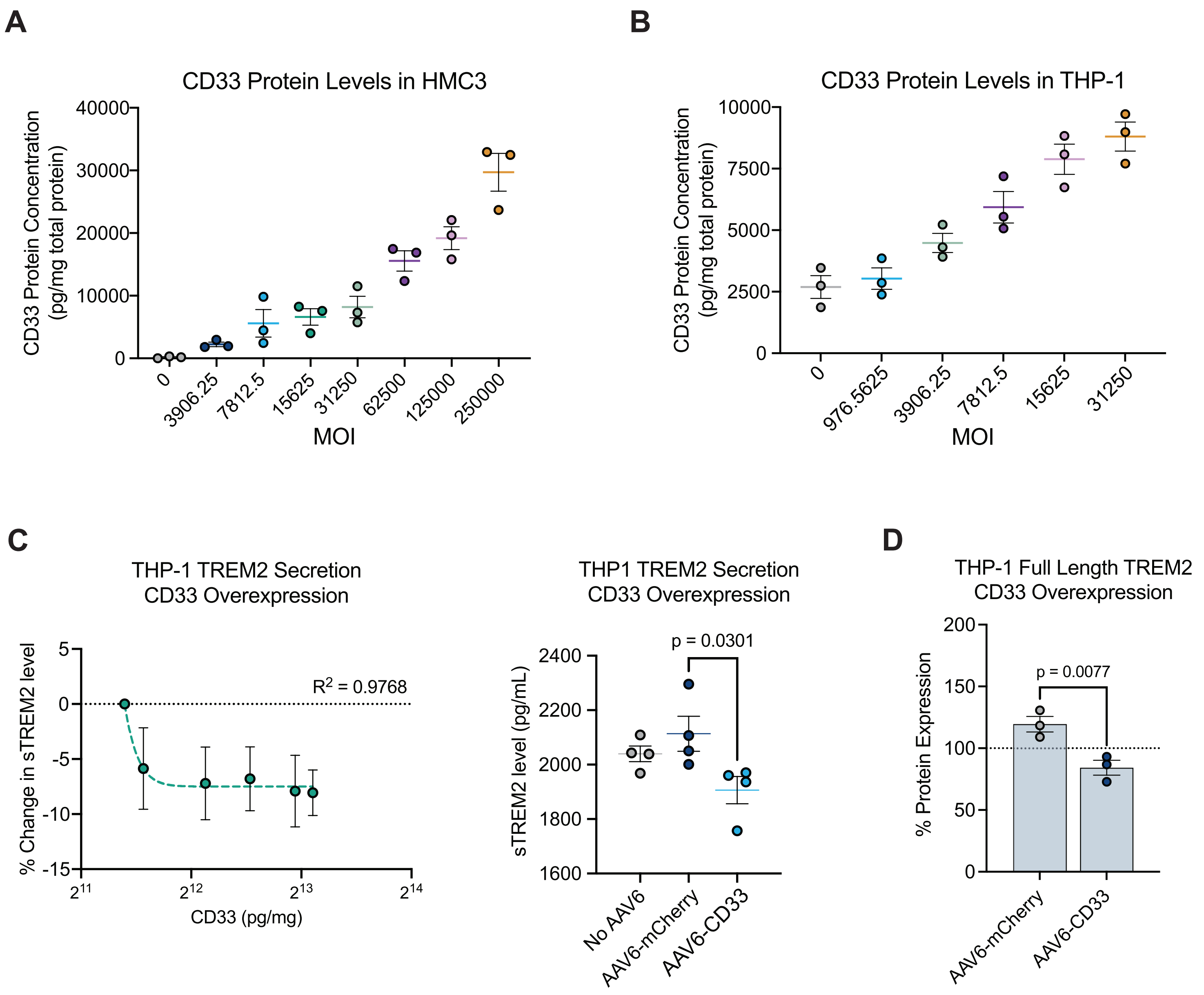

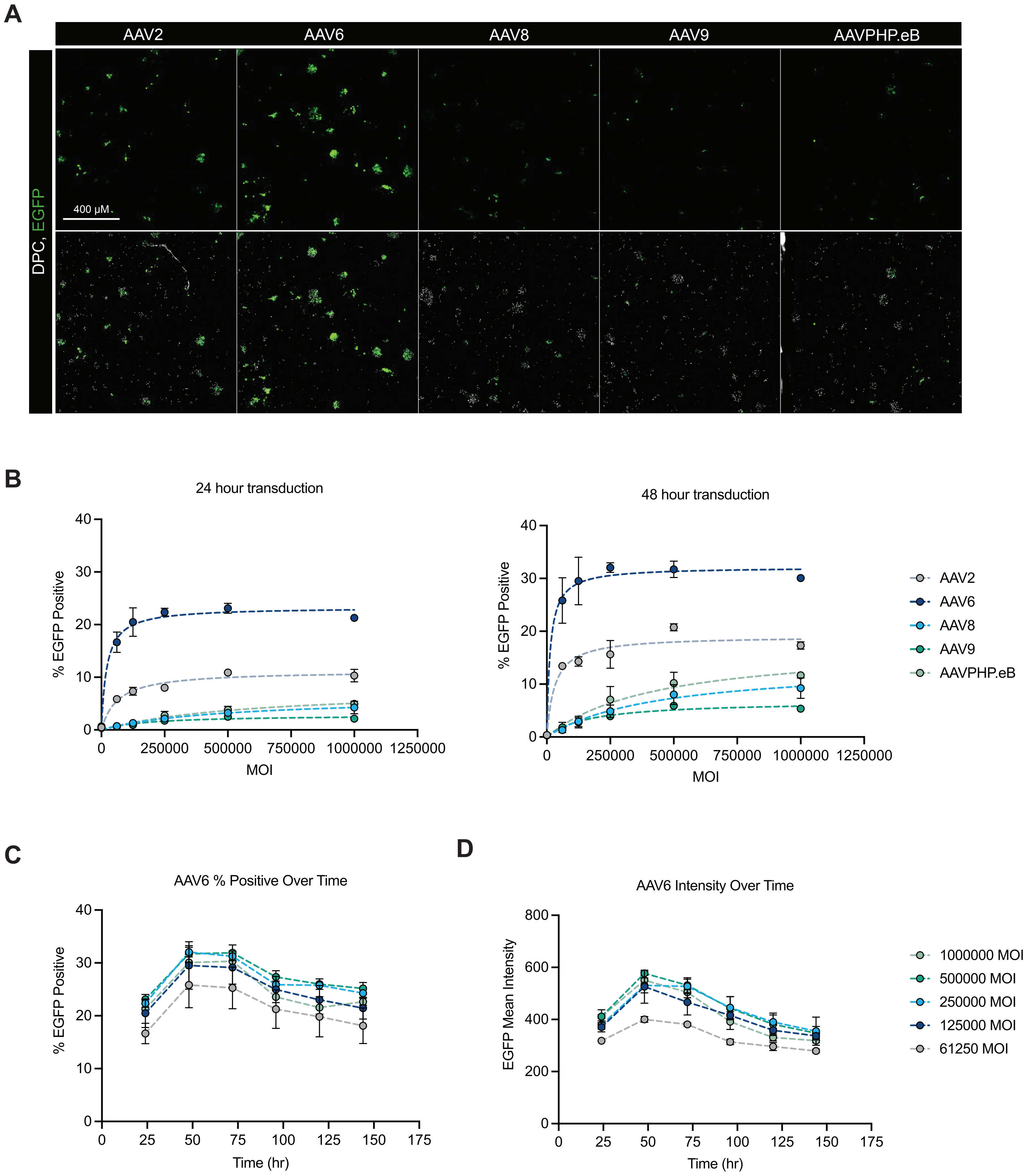
